# DJ-1 (Park7) affects the gut microbiome, metabolites and development of Innate Lymphoid cells (ILCs)

**DOI:** 10.1101/776005

**Authors:** Yogesh Singh, Christoph Trautwein, Achal Dhariwal, Madhuri S Salker, Mohammed Alauddin, Laimdota Zigmare, Lisan Pelzl, Martina Feger, Jakob Matthes, Nicolas Casadei, Michael Föller, Vivek Pachauri, David S Park, Tak W Mak, Julia S Frick, Diethelm Wallwiener, Sara Y Brucker, Florian Lang, Olaf Riess

**Author notes:** senior authors (Florian Lang, Olaf Riess). Address for correspondence Yogesh Singh, PhD, Institute of Medical Genetics and Applied Genomics, Calwerstraße 7, 72076, Tübingen University, Tübingen, Germany.

## Abstract

The proper communication between gut and brain is pivotal for maintenance of health and dysregulation of the gut-brain axis can lead to several clinical disorders. Also, in Parkinson’s disease (PD) 85% of all patients experienced constipation long before showing any signs of motor phenotypes. For differential diagnosis and when it comes to preventive treatment there is an urgent need for the identification of biomarkers indicating early disease stages long before the disease phenotype manifests. DJ-1 is a chaperon protein involved in the protection against PD and genetic mutations in this protein have been shown to cause familial PD. However, how the deficiency of DJ-1 modifies the PD risk remains incompletely understood. In the present study we provide evidence that DJ-1 is implicated in shaping the gut microbiome including their metabolite production or innate immune cells (ILCs) development. We revealed that in 4 months old mice genetic deficiency of DJ-1 leads to significantly decrease in several bacterial genera and significantly increase in two specific genera, namely *Alistipes* and *Rikenella*. DJ-1 deficient mice have a higher production of calprotectin/MCP-1 inflammatory protein - a known protein involved in colonic inflammation – and significantly higher expression of glial fibrillary acidic protein (GFAP) than control littermates. Expression of a-Synuclein, a key protein in Lewy bodies, in the colon was not significantly different between genotypes. Metabolic profiles of feces extracts analysed by H^1^-NMR spectroscopy showed increased short chain fatty acids (SCFAs) and decreased amino acid levels, suggesting a general switch from protein towards fibre degrading strains in DJ-1 deficient mice. We observed that Malonate - which is known to influence the immune system – has significantly higher concentrations in DJ-1 deficient mice. Moreover, DJ-1 deficient mice have high levels of the phenol derivate 3-(3-Hydroxyphenyl) propanoic acid (3-HPPA) which is a breakdown product of aromatic substrates like tyrosine, phenylalanine and polyphenols. DJ-1 deficient mice also showed significantly reduced percentage of ILCs. Thus, our data suggests that absence of DJ-1 leads to increase in gut inflammatory bacteria composition, deregulated metabolites and dysregulated innate immunity which could be a key factor in the initiation of PD disease in the gut, and potentially also in brain during disease progression.

## Introduction

Parkinson’s disease (PD) is the most common movement disorder and the second most prevalent neurodegenerative disease in humans (*1*). Clinically, PD patients suffer from resting tremor, rigidity, bradykinesia and altered gait (*1*). PD is an incurable neurodegenerative disease distinguished by the loss of neurons predominantly in the *substantia nigra pars compacta* (*SNpc*) region in the brain and the presence of Lewy bodies in the surviving neurons, however the exact cause of PD and how the disease process is triggered remained incompletely understood (*2*). Most of the PD cases are sporadic, however, several rare genetic forms of the disease have been identified that have contributed prominently to our understanding of the mechanisms underlying disease pathogenesis (*3*). In the Lewy bodies, presence of these intracellular protein inclusions is mainly comprised of misfolded alpha-Synuclein (α-Syn), which has also been shown to be genetically linked to familial and sporadic forms of PD (*4, 5*).

In addition to α-Syn-related genetic links with PD, several other mutations such as PARK7 (DJ-1), Parkin, UCH-L1, Pink1 and dardarin genes account for sporadic cases with early-onset recessive PD (*6, 7*). DJ-1 is a small ubiquitously expressed protein implicated in several pathways associated with PD pathogenesis (*3*). DJ-1 protein is encoded by the PARK7 gene and comprises of 189 amino acid long (*8*). DJ-1 is localized primarily in the cytoplasm, however it can also be found in the nucleus and linked with mitochondria in the cell (*8*). DJ-1 is involved in several cellular functions, serving as an oxidative stress sensor (via a cysteine residue at position 106, C106), a protein chaperone, a protease, an RNA-binding protein, a transcription regulator, a regulator of mitochondria function and a regulator of autophagy (*9*). It is still not clear which of these processes are responsible for DJ-1-dependent pathogenesis in PD.

Previous studies revealed that DJ-1 is involved in the modulation, aggregation and toxicity of α-Syn (*8, 10–12*). Double transgenic mice for DJ-1 deficiency and α-Syn expression (expressing pathogenic Ala53Thr human a-Syn) called M83-DJ-1 null mice, revealed that onset of disease and pathological changes were not different when compared with single transgenic M83 mice line and concluded that α-Syn and DJ-1 mutation may lead to PD *via* independent mechanisms *(13)*. However, a recent *in vitro* study revealed that DJ-1 directly binds monomeric/oligomeric α-Syn and showed that DJ-1 interacts with α-Syn in living cells (*8*). Nevertheless, overexpression of DJ-1 protects against neurodegeneration in the yeast and Drosophila models *in vivo* (*8*). DA neurons number in the *SNpc* region and fibre densities and dopamine levels in the striatum were reported to be normal in DJ-1^−/−^ mice. Enhanced striatal denervation and dopaminergic neuron loss was induced by 1-methy-4-phenyl-1,2,3,6-tetrahydropyridine (MPTP) together with amphetamine in DJ-1^−/−^ mice (*7*). Additionally, DJ-1^−/−^ mice, which were backcrossed with C57BL/6 mice (known as DJ-1C57^−/−^) suffered from early-onset unilateral loss of dopaminergic (DA) neurons in their *SNpc*, progressing to bilateral degeneration of the nigrostriatal axis with aging as well as mild motor behaviour deficits at aged time points (*14*). The authors suggested that the DJ1-C57 model effectively recapitulates the early stages of PD and allowing to study the preclinical aspects of neurodegeneration (*14*). Our own studies with DJ-1^−/−^ mice models also suggested that DJ-1 protein is also involved in the maintenance the physiology of adaptive immune CD4^+^ T cells and their development and functions by regulating the sodium hydrogen exchanger 1 (NHE1) and ROS formation (*15, 16*). Keeping in mind that different DJ-1^−/−^ mice lines with different background could prone to develop neurodegeneration and some are resistance then it is possible that in addition to genetic changes additional environment and unknown factors may play a crucial role for the disease development in DJ-1^−/−^ mice. One of factor could be the gut microbiome which could be governing the resistant phenotype. Thus, studying the role of gut microbiome in the context of disease development in DJ-1^−/−^ mice model is warranted to understand the PD pathophysiology to develop novel tools for pre-clinical studies.

Trillions of bacteria live and reside in the gut named jointly as the gut microbiome. They are important for normal functioning of the intestine (*17*). Recent advances in metagenomics techniques leads to step into this new world of research to uncover the profound impacts that the microbiota may have on neurodevelopment and diseased of the central nervous system (CNS) (*18, 19*). An essential function of the gastrointestinal tract (GIT) is to perceive and react to external signals such as environment, food, and xenobiotics (*20*). Studies from germ-free mice (GF) and antibiotic-treated mice suggested that bacteria are pivotal in hippocampal neurogenesis maintenance as well as spatial and object recognition (*21*). In mice, antibiotics treatment changed transiently the microbiota, and as a result increased expression of the brain-derived neurotropic factor (BDNF) in the hippocampus and enhanced exploratory behavior was observed (*22*). Further studies suggested that the microbiota promotes enteric and circulating serotonin (5-hydroxytryptamine, 5-HT) production from colonic enterochromaffin cells, modulates the GIT mobility, platelet functions (*23*), affects anxiety, hyperactivity, and cognition (*24, 25*). Dysbiosis (alterations to the microbial composition) of the human microbiome has not only been described in mice, but also in persons with several neurological diseases (*26*). For example, fecal and mucosa-associated gut microbes are different between individuals with PD and healthy controls (*27–31*). Thy1-α-Syn [ASO] transgenic PD mouse model study also suggested that PD derived microbiota have adverse effect on the Parkinson’s pathogenesis including the accumulation of α-Syn and change in the motor phenotype (*32*). Several other mouse studies also suggested the gut dysbiosis in chemically induced toxins and other PD models (*32−38*). Human studies from fecal metabolites suggested that PD patients have reduced short chain fatty acids (SCFAs), which are the metabolic products of certain gut bacteria (*31*). However, how bacterial produced gut metabolites are affecting the neurodegenerative diseases are not understood and an area of active research.

Immune cells are capable of engaging in direct communication with enteric neurons (*18, 20*). The extent of the functional impact of neuro-immune synapses is not clear yet however published studies advocated that activated immune cells can temper neuronal activity *via* the release of neurotransmitters, metabolites and cytokines (*19, 39, 40*). Based on the common occurrence of GIT symptoms in PD, dysbiosis among PD patients, and evidence that the microbiota impacts CNS function, we hypothesized that DJ-1 protein could also be involved in the regulation of the gut microbiome and inflammation. Herein, we report that the microbiota composition is dysregulated, change in the innate immunity, increased inflammation in the colon and feces and dyregulated metabolites in the absence of DJ-1 in young adult mice.

## Results

### Gut microbiome dysbiosis (dysregulation of intestinal bacterial community signatures) in young DJ-1^−/−^ mice

Recent studies in PD patients have described the potential link with gut microbial abundance and Parkinson’s pathogenesis (*28, 29, 31*). Further findings suggested that the brain-gut axis interactions are controlled by the gut microbiome through immunological, neuroendocrine and direct neural mechanisms, respectively (*19, 39*). Therefore, a clear understanding of the microbiota-gut-brain axis interaction could bring a new insights in the pathophysiology of PD and allow an earlier diagnosis with a focus on peripheral biomarkers within the enteric nervous system (*41*). However, how the DJ-1 deficiency has any potential effects on the gut microbiome is not known yet. To understand this process in more detail, we used 16S rRNA sequencing method and characterized the gut microbiome from DJ-1 deficient (DJ-1 KO or DJ-1^−/−^) and control littermate wild-type (WT) fecal samples from 4 months old animals. Both the WT and DJ-1^−/−^, were kept in the same cage and animals were kept in 5-6 different cages to nullify the cage effect as mice are coprophagic in nature. Heterozygous mothers were mated to obtain DJ-1^−/−^ and WT (DJ-1^+/+^) animals to minimize the effect of the maternal microbiome. Data analysis of 16S rRNA sequencing reads were performed using the *MEGAN-CE* microbiome analyzer software (*42*) as well as *MicrobiomeAnalyst* tool: a web-based tool for comprehensive statistical, visual and meta-analysis of microbiome data (*43*) as described in the materials and methods sections in detail.

Total number of reads were significantly less in DJ-1^−/−^ samples compared with WT control littermate samples (Fig. 1a). Our microbiome data at phylum level analysis suggested that 4 months old DJ-1 deficient mice tended to have higher alpha diversity Chao 1 and lower Shannon-Weaver index: species richness within a single microbial ecosystem (*44*) of gut microbiome compared with the control littermate WT, but it did not reach significance level (Fig. 1b, c). Similarly, beta diversity: diversity in microbial community between different environments (*44*) tended to be reduced (UniFrac: both weighted and Unweighted), a difference, however, again not reaching statistical significance. DJ-1^−/−^ mice have significantly higher abundance of *Bacteroidetes* and significantly less abundance of *Firmicutes* and *Cyanobacteria* compared with WT in an aged matched control littermate (Fig. 1d). Further, when we mined the data for the overall composition of the gut bacterium at the phylum level, we found that indeed several bacterial phyla tended to be different, but the difference again did not reach statistical significance (Fig. 1e). As earlier studies suggested that Firmicutes/Bacteroidetes (F/B) could help to predict the functionality of the microbiome, hence, we calculated the F/B ratio and found that in DJ-1^−/−^ mice the F/B ratio was statistically significantly decreased when it was compared with control WT littermates (Fig. 1e).

**Fig. 1:**
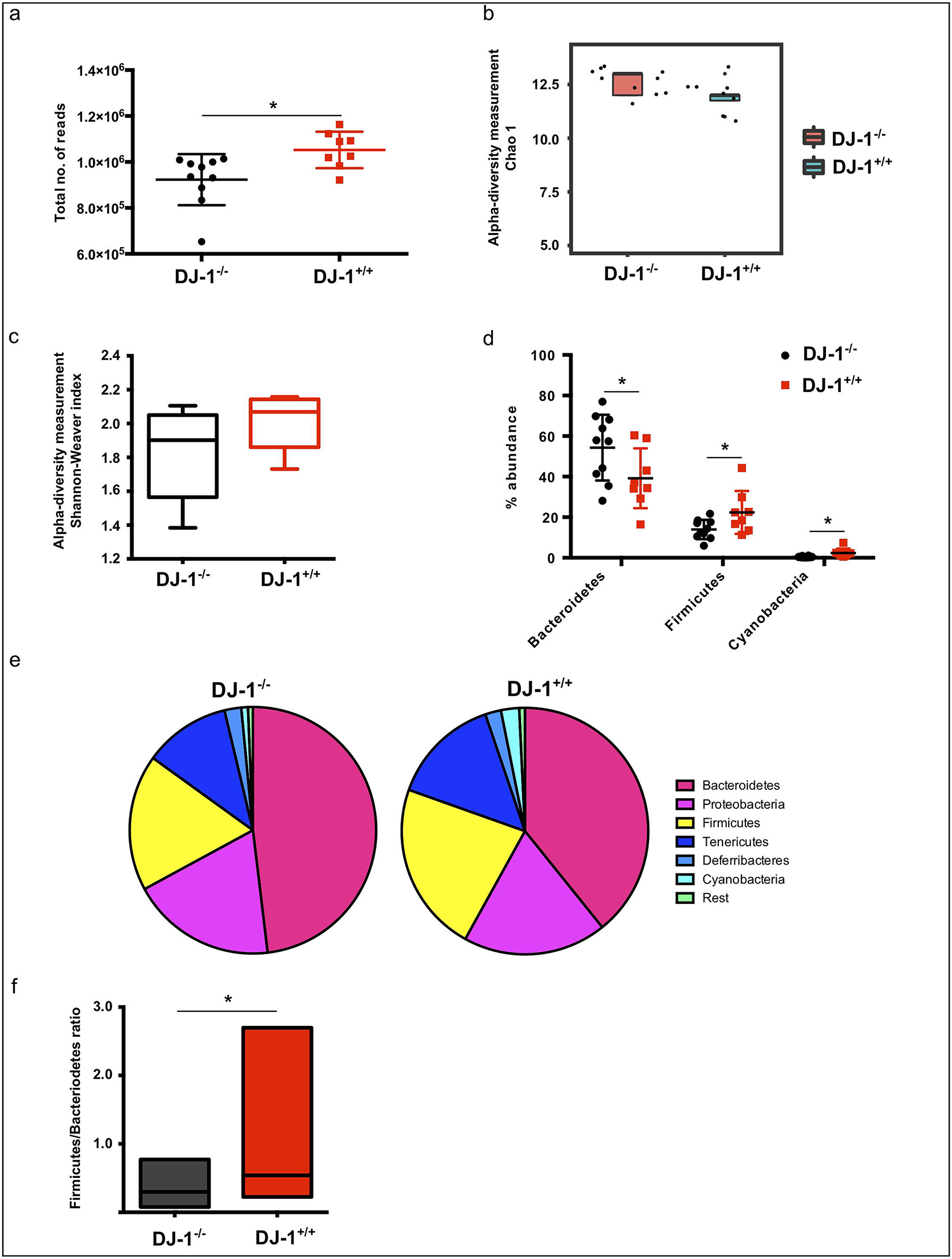
Gut dysbiosis is prevalent in four months old DJ-1^−/−^ mice. (a) Total number of sequences read obtained from sequencing of colon fecal samples from DJ-1^+/+^ (n=8) and DJ-1^−/−^ (n=10) mice. (b) Measurement of alpha diversities (Chao1 and Shannon-Weaver index) for WT and DJ-1^−/−^ mice. (c) Bacteroidetes, Firmicutes and Cyanobacteria were significantly different between WT and DJ-1^−/−^ mice. (d) Bacterial abundance data (mean±SD) presentation at phylum level in WT and DJ-1^−/−^ mice. (e) Firmicutes/Bacteroidetes ratio in WT and DJ-1^−/−^ mice, it was significantly reduced in DJ-1^−/−^ mice compared with WT. (f) Firmiuctes/Bacteroiidetes (F/B) ratio was decreased in DJ-1^−/−^ compared with WT mice. *represents the p value of <0.05 using Student’s unpaired t-test.

We further calculated the alpha and beta diversities at the genera level and found that both the alpha diversity (Chao1 and Shannon-Weaver index) was significantly reduced in DJ-1^−/−^ mice compared with control littermate WT animals (Fig. 2a, b). Furthermore, analysis using Bray-Curtis principal component analysis (PCoA) with MEGAN-CE and UniFrac to differentiate between the two different mice lines (WT and DJ-1^−/−^) and found that indeed, at a younger age both the animals were different in their bacteria abundance and clustering. Beta diversity was also calculated using MicrobiomeAnalyst tool (*43*) (PERMANOVA). As a result, WT and DJ-1^−/−^ were different at both at the genus (p value=0.03) and species level (p value=0.006) respectively. More than 1% bacterial genera were represented in a pie chart and clustering of bacterial genera were shown as heat map (Fig. 2c, d). In total 17 bacterial genera were significantly changed in the DJ-1 deficient animals (Fig. 2e). Most of the bacterial genera were significantly downregulated (*Anaerotruncus, Comamonadaceae, Acholeplasma, Streptococcus, Blautia, Ruminococcaceae, Cyanobacteria, Bilophila, Ruminococcaceae-uncultured, Peptococcus, Merismopedia, Oscillospira, Pseudobacteroides, Anaerosporobacter,* and *Robinsoniella*), only the opportunistic symbionts *Rikenlla* and *Alistipes* were significantly upregulated (Fig. 2e). The abundance analysis at species levels confirmed that both the pathogenic bacteria *Alistipes* sp and *Rikenella sp* were significantly higher in DJ-1 deficient mice with the WT littermate controls (Fig. 3a, b). Although 16S rRNA sequencing is not specific enough to characterize the bacterium at species/strain level, however, it is sensitive enough to get an approximate idea which species could be present. Henceforth, we mined our 16S rRNA data and found that indeed both unknown *Rikenella sp.* and *Alistipes timonesis* were significantly upregulated. The bacterial *Prevotella sp* was also significantly more abundant in DJ-1^−/−^ mice.

**Fig. 2:**
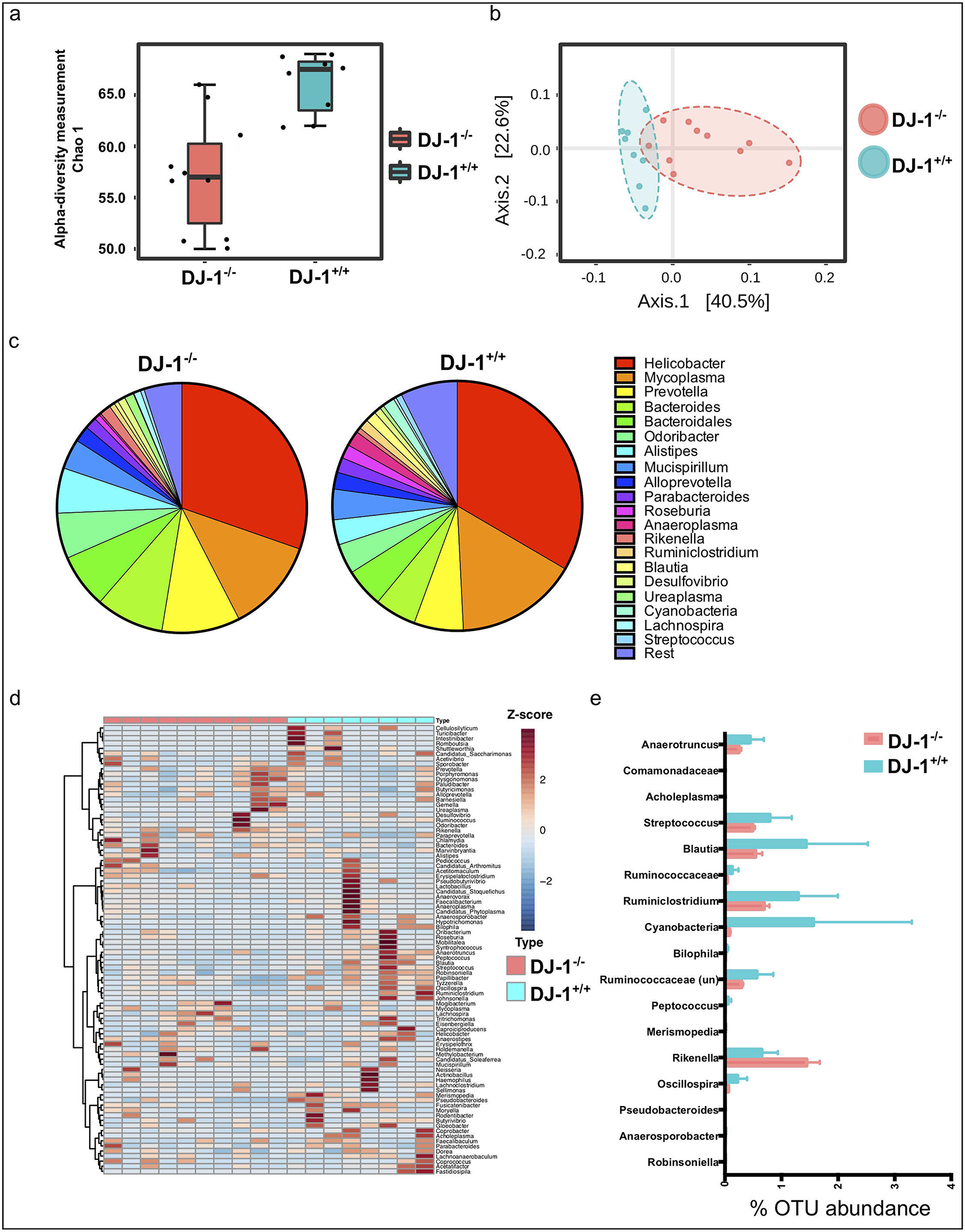
Increase in *Rikenella* and *Alistipes sp.* bacteria in DJ-1^−/−^ mice compared with WT at four months of age. (a) Measurement of alpha (Chao1) diversity in WT and DJ-1^−/−^ mice. (b) Beta diversity (UniFrac) for WT (n=8) and DJ-1^−/−^ (n=10) mice. Both WT and DJ-1^−/−^ mice clusters in a separate group. (c) A representative image of abundance of bacteria at genus levels in all WT (n=8) and DJ-1^−/−^ (n=10) mice fecal samples. (d) Heat map represents the bacterial abundance in WT and DJ-1^−/−^ mice in all the animals. The abundance of bacteria at genera level are represented in Z-score. (e) Average statistically significant bacterial abundance at genus level (only significantly abundant genera are shown). *represents the p value of <0.05 using Student’s unpaired t-test.

**Fig. 3:**
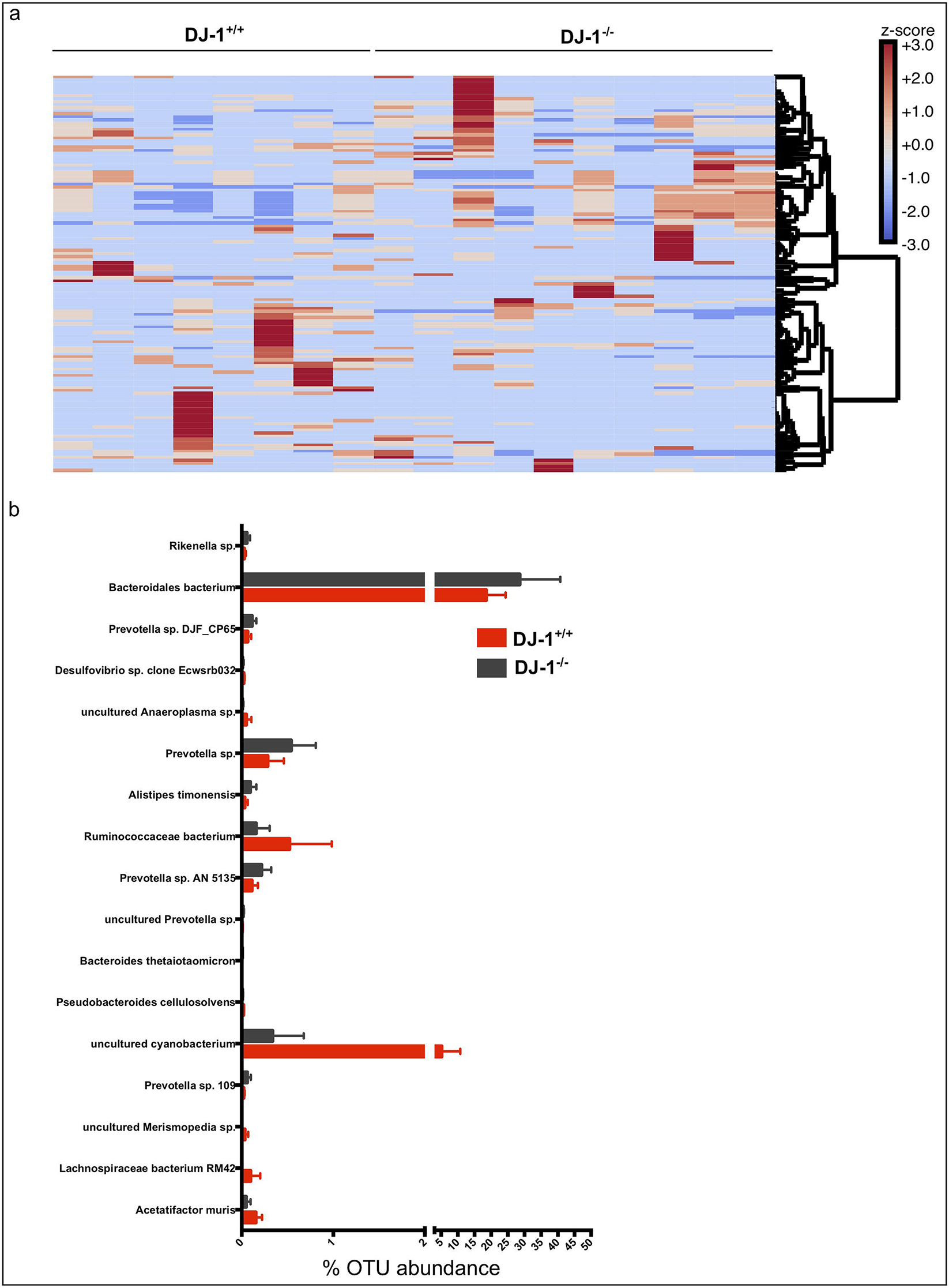
Significant increase in *Rikenella* sp. and *Alistipes timonensis* in DJ-1^−/−^ mice compared with WT. (a) Clustering of all bacteria at species level. (b) Bar diagram (mean±SD) shows all the significantly different bacteria at species level in WT and DJ-1^−/−^ mice. *represents the p value of <0.05 using Student’s unpaired t-test.

We next used the functional gene prediction profiling through Tax4Fun from our 16s rRNA data set (*45*) and output of these functional profile at KEGG metabolism level was visualized in SDP module of MicrobiomeAnalyst tool (*43*). The association analysis was performed using Global test algorithm between the two groups (DJ-1^−/−^ and DJ-1^+/+^ mice). We observed that most of genes were predominantly associated with the pathways of amino acid, carbohydrate, energy, vitamins, cofactors and nucleotides metabolisms in WT and DJ-1 deficient mice (Suppl. Fig. 1). Overall, our data of the 16S rRNA bacterial sequencing analysis suggested that genetic deficiency of DJ-1 affects the bacterial composition and functions in the gut, even before these animals develop any disease phenotype.

### Increased inflammation in the feces and colonic tissues, compromised innate immune system in DJ-1^−/−^ mice

The dysregulated immune system could lead to inflammation and increased permeability of the gut (*46, 47*). Thus, we further characterized the inflammatory protein calprotectin in the feces. Due to leukocytes shedding in the intestinal lumen, pro-inflammatory proteins such as calprotectin (S100A8/S100A9) can be detected and measured in the stool by ELISA (*48*). The concentration of calprotectin is directly proportional to the intensity of the neutrophil infiltrate in the gut mucosa. Calprotectin is released from neutrophils, monocytes, macrophages and epithelial cells in the case of the gastrointestinal tract inflammation (*48*); another study suggested that PD patients have higher expression of fecal calprotectin and zonulin (marker of increased gut permeability) proteins (*47*) and other study also suggested enhanced inflammation in PD patients (*49*). To validate whether increased gut dysbiosis in DJ-1^−/−^ mice have higher inflammation in the gut or not, we measured the calprotectin levels. In DJ-1^−/−^ mice calprotectin appeared to be higher than in WT, a difference, however, not reaching statistical significance (Fig. 4a). Moreover, we also measured other inflammatory cytokines in the feces and found that the monocyte chemotactic protein-1 (MCP-1) was significantly upregulated in DJ-1 deficient mice compared with littermate WT control (p =0.034) (Fig. 4b). Other pro-inflammatory cytokines (IL-12p70, TNF-α, IL-17A, IFN-γ, IL-23 and IL-6) were tended to be upregulated in the feces from DJ-1^−/−^ compared with WT mice, however, did not reach at significance level (Suppl. Fig. 2). The cytokines such as GM-CSF, IFN-β and IL-27 were significantly down regulated in DJ-1^−/−^ mice compared to WT (Supp. Fig. 2). Additionally, previous studied suggested that pro-inflammatory cytokines are able to increase glial fibrillary acidic protein (GFAP) expression in enteric glia (*50*) and PD patients have also enhanced inflammation (*49*), therefore, we explored the quantification of GFAP from the colon tissue. Our Immunoblotting data suggested that DJ-1^−/−^ colon have significantly higher GFAP protein compared with WT control littermate mice (Fig. 4c).

**Fig. 4:**
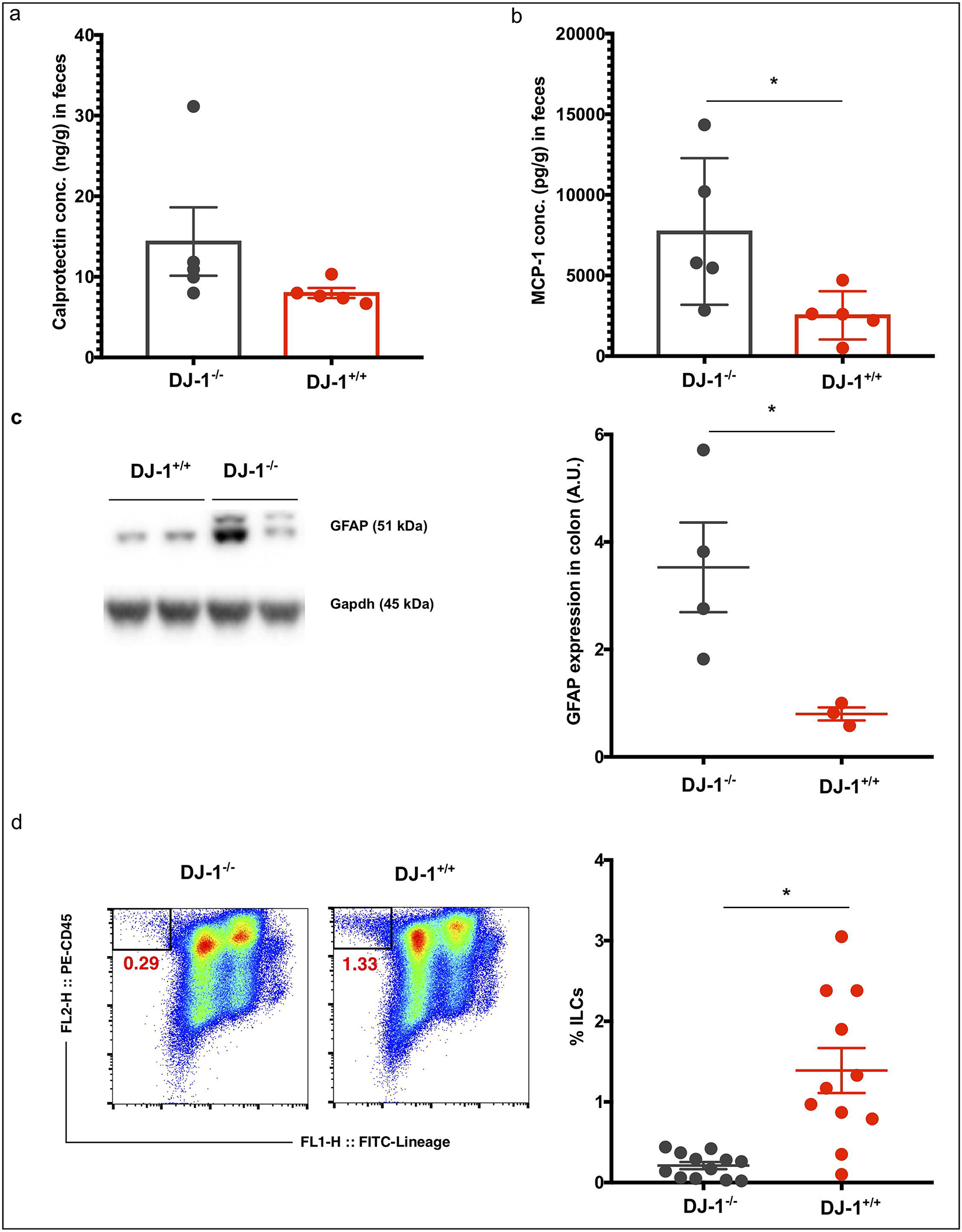
Increased inflammation and decreased ILCs in DJ-1^−/−^ mice compared with WT. (a) Calprotectin level (mean±SEM) measured by ELISA in WT and DJ-1^−/−^ mice fecal samples (n=5). (b) DJ-1^−/−^ mice have significantly higher production of the MCP-1 (mean±SEM) cytokine measurement from the feces. p values (0.04 Student’s t-test). * represents the p value of <0.05. (c) Original Immunoblots from GFAP expression and quantification of the protein expression. Bar diagram (mean±SEM) shows the statistical significantly higher expression of GFAP in the colon tissues (p value =0.03) of the DJ-1^−/−^ mice compared to control WT mice. (d) Left hand side figure shows representative FACS plots from WT (n=8) and DJ-1^−/−^ (n=10) mice spleen samples. X-axis represents Lineage markers of lymphoid cells (CD3, CD5, CD3, CD11b, CD11c, F4/80, Gr-1, B220 or CD19, and Ter119) and y-axis represents lymphoid marker CD45. Right hand side Bar diagram (mean±SEM) shows significantly lower in DJ-1^−/−^ (n=12) compared with WT (n=11). *represents the p value of <0.05 using Student’s unpaired t-test.

Unwanted activation of the innate immune system from small intestinal bacterial overgrowth leads to increased intestinal permeability and may cause systemic inflammation (*41*). In the same perspective, unwanted activation of enteric neurons and enteric glial cells could follow the initiation of α-Syn accumulation and misfolding in the gut and the brain (*51*). In addition to the innate immune system, the adaptive immune system could also be affected and changed by bacterial proteins cross-reacting with host antigens and could lead to severe inflammation of the host tissues (*52*). Elevated α-Syn expression impairs innate immune cells function (*53*). Our previous studies suggested that DJ-1^−/−^ mice have less induced regulatory T cells (Tregs) (*16*). We speculated that the defect could be in the adaptive immune cell development or functions in the DJ-1^−/−^ mice. However, involvement of innate lymphoid cells (ILCs) in the pathogenesis of PD has not been described yet. To delineate the role of ILCs, we first characterized these cells using Flow cytometry and found that CD45^+^Lineage^−^(CD3, CD5, CD3, CD11b, CD11c, F4/80, Gr-1, B220 or CD19, and Ter119) were significantly less abundant in DJ-1^−/−^ mice compared with WT control littermates in the spleen (Fig. 4d).

### Bacterial metabolites in the feces of DJ-1^−/−^ mice are dysregulated

Our results suggested that DJ-1^−/−^ mice suffer from gut dysbiosis and as a result could have differences in their fecal metabolite production resulting in changed physiology and disease outcome. Several previous studies have identified that gut bacteria control different metabolites which are involved in neurodegenerative diseases (*19, 54–56*). Therefore, we used ^1^H-NMR based metabolomics for the identification and quantification of fecal metabolites as described elsewhere in detail (*57, 58*). The detected compounds cover a comprehensive range of metabolite classes such as amino acids, SCFAs, phenols, amines, carbohydrates, purines, alcohols and others.

Our extraction procedure yielded very rich and high-quality ^1^H-NMR spectra with final TSP linewidths < 1 Hz. A total of 40 metabolites could be annotated and quantified in all samples, mainly amino acids, short chain fatty acids, carbohydrates and nucleotides. The clustering and non-clustering of all metabolites is shown in the heatmap (Supp. Fig. 3). Aside from known metabolites we identified in half of the samples high levels of 3-(3-Hydroxyphenyl)propanoic acid (3-HPPA) (CAS 621-54-5), a phenol derivative which has been shown to be able to readily cross the gut epithelium (*59*) into the blood and brain (*60*). 3-HPPA has been shown to be formed mainly by *Clostridium, Escherichia and Eubacteria* species (*61, 62*).

Using classical volcano plot analysis with a fold change threshold of 1.5 and student’s t-test we identified 9 metabolites as highly discriminating the two groups (Fig. 5a) with Malonate (p = 0.006) being top for DJ-1^−/−^ and Valine (p = 0.008) for WT (Fig. 5b). Further multivariate statistics using orthogonal T-score (Partial Least Squares Discriminant Analysis: oPLS-DA) was used to find out the similarity between DJ-1^−/−^ and WT feces samples (Fig. 6a,b). Here, we found a strong separation of both groups and the corresponding s-plot demonstrates that SCFAs and amino acids are the most discriminating metabolites for clustering. Further, we applied variable importance of projection (VIP) PLS-DA analysis to identify correlation patterns and found again high significance of SCFAs, amino acids as well as carbohydrates and aromatic compounds metabolism with VIPs-scores > 1 (Fig. 6c). Interestingly the high abundance of SCFAs in DJ-1^−/−^ shows a high correlation with the newly identified metabolite 3-HPPA (Fig. 6d and Suppl. Fig. 4). Thus, overall data suggested that DJ-1^−/−^ mice have impaired protein degradation -low levels of amino acids while fiber digestion was enhanced - high SCFAs production and could potentially trigger immune response and gut epithelial dysbiosis.

**Fig. 5:**
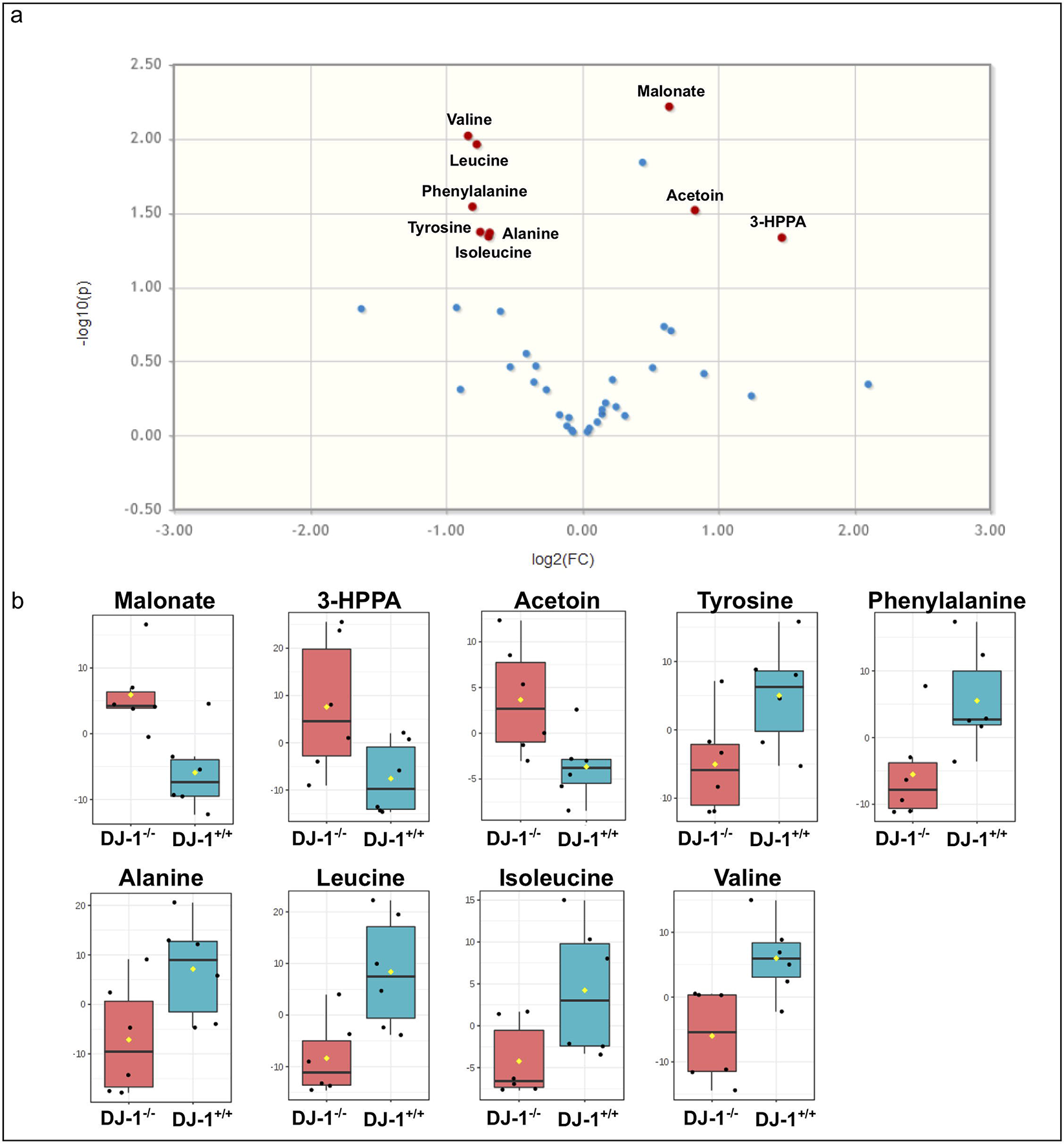
Reduced amino acids and increased SCFAs in feces from DJ-1^−/−^ mice feces. (a)Volcano plot analysis with boxplots of quantitative metabolite data from ^1^H-NMR spectroscopy. (b) A total of nine metabolites showed p-values < 0.05 and fold changes > 1.5 between the two groups with Valine (high in WT) and Malonate (high in DJ-1^−/−^) being the most significant ones.

**Fig. 6:**
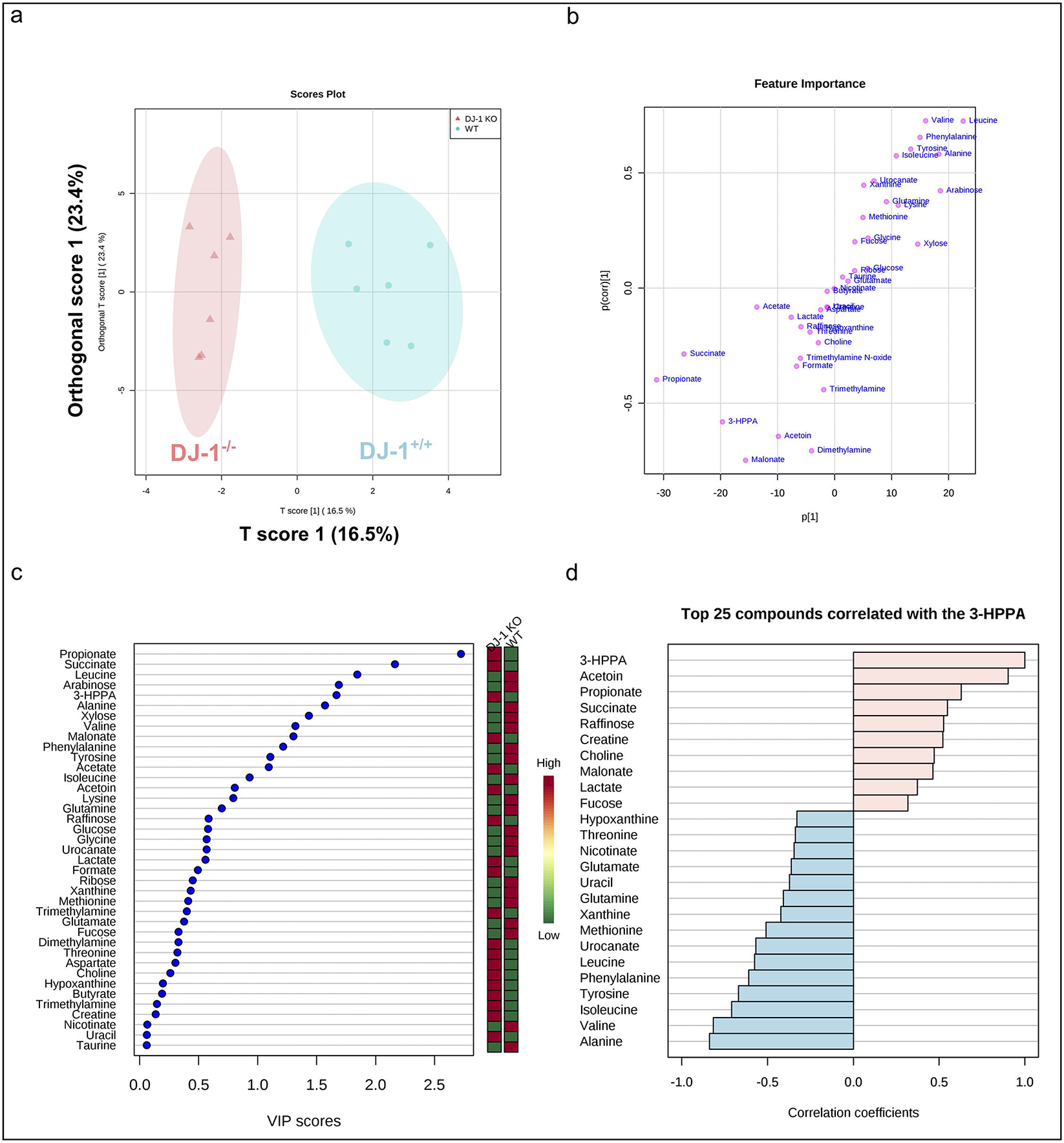
DJ-1^−/−^ feces metabolites differently clusters and have difference in metabolite production than the WT feces. (a,b) Orthogonal partial least squares discriminant analysis (oPLS-DA) with the corresponding s-plot clearly separates WT and DJ-1^−/−^ groups with short chain fatty acids and amino acids being the most discriminating metabolites, indicating a general switch from protein towards fibre degrading strains in DJ-1^−/−^ feces. (c,b) Partial least squares discriminant analysis PLS-DA with VIP (variable importance in projection) scores for the top 20 metabolites and pattern hunter of the newly in feces identified phenolic metabolite 3-(3-Hydroxyphenyl) propanoic acid.

## Discussion

Over the past years, our understanding of human-associated microorganisms has been vastly improved beyond that of a certain species toward an appreciation of the diverse and niche-specialized microbial communities that develop in the human host (*63*). Our intestinal tract especially the colon contains a largest reservoir of different microorganisms such as bacteria, viruses, fungi etc. interacting with the host epithelial and immune cells. These microorganisms orchestrate the primary and secondary metabolism on the gut mucosal surface and influence the other body organs, mucosal and hematopoietic immune functions (*63*). Thus, it is not surprising that modulation in the composition and function of the gut bacteria has been linked with several chronic diseases including gastrointestinal inflammation, metabolic disorders as well as cardiovascular and neurological diseases (*41, 64–66*). Recent discoveries in the field of the gut microbiome and their role in the neurodegeneration in patients and animal models highlights an important link among each other (*67*). As most of the common neurodegenerative diseases such as Alzheimer’s disease and PD occurs late in the life (*40, 68*) biomarkers allowing early diagnosis would be invaluable.

### Bacterial composition and dysbiosis

Animal models are useful tools to understand the pathophysiological functions of disease as these models allow us to manipulate the course of the disease (*69*). To understand the impact of the gut microbiome in the pathophysiology of PD in early life prior to development of any phenotype or symptoms, we used DJ-1^−/−^ animal models. The DJ-1 mutation has been found in 1-2% early-onset recessive PD and PD patients containing DJ-1 mutation are normally responsive to levodopa (*8*). We found that two novel bacteria such as *Alistipes* and *Rikenella* genera were highly abundant in DJ-1^−/−^ mice compared with WT control littermate animals. Both the alpha and beta diversity analysis suggested that both genotypes of animals have a distinct microbiota diversity and community. *Alistipes* and *Rikenella sp* have been described earlier to be involved in the inflammatory bowel disease as well as colon cancer development (*70*). In humans (colorectal cancer patients) a number of *Bacteroides* and *Parabacteroides* species, along with *Alistipes putredinis, Bilophila wadsworthia, Lachnospiraceae bacterium* and *Escherichia coli* were enriched in carcinoma samples compared with both healthy and advanced adenoma samples (*71*). Gut commensals such as *Bifidobactium animalis* and *Streptococcus thermophilus*, on the other hand, decreased in faeces from adenoma or carcinoma patients, consistent with deviation from a healthy microbiome (*71*). *Alistipes* and *Ruminococcus* are shown to be positively correlated with TNF-α production after anti-IL-10R/CpG oligonucleotide immunotherapy in C57BL/6 mice suffering from MC38 colon carcinoma (*72*).

Chondroitin Sulfate has been shown to increase the abundance of *Rikenella*, a genus of sulphate reduced bacteria, but did not significantly change the abundance of *Akkermansia muciniphila (73)*. Surprisingly, *Rikenella* was exterminated by cephalosporin whereas *Rikenella* blossomed on exposure to berberine. However, *A. muciniphila* was eliminated by this antibiotic compound. Cephalosporin significantly reduced colonic mucus lesions and delayed the early pathogenesis of dementia, steatohepatitis, and atherosclerosis. Berberine significantly aggravated colonic mucus lesions and enhanced multi-systemic pathogenesis (*73*). Thus, *Rikenella* could be involved in sulphate reduction in DJ-1^−/−^ mice and could be a triggering factor to induce colonic mucus lesions and PD disease progression.

### Intestinal inflammation regulation by inflammatory bacteria and immune cells

Intestinal inflammation has been linked with gut permeability and induction of immune response as well as function of gut neurons (*19, 63*). LPS challenged DJ-1^−/−^ mice IFN-γ and interferon-inducible T-cell α chemoattractant were enhanced in the *SNpc* and microglial cells compared with WT animals (*74*). These results suggested that interaction of genetic defect with inflammatory mediators could participate in the development of PD. Our data also suggested that DJ-1^−/−^ mice suffer from higher fecal inflammation as well as neuronal inflammation (GFAP), which could be linked with neurodegeneration. However, the DJ-1^−/−^ mice do not develop any phenotype in unchallenged conditions. It is worth to note that the study was performed at early stage before any neurological symptom develops in the brain. Recent studies suggested that DJ-1C57^−/−^ with a different genetic background (it is backcrossed 14 generations with C57BL/6 mice), is more penetrant in disease phenotype development, as these animals develop the phenotype within 3 months of age and progress to by 12 months (*14*). The exact mechanism is unknown, but it could possibly be correlated with the genetic backgroung composition in an interplay with the gut microbiome development. However, further studies, in particularly also of the microbiome, are warranted to understand the disease penetration in DJ-1C57^−/−^ mice and DJ-1^−/−^ animals. DJ-1^−/−^ mice have no appreciable change in α-Syn expression in the colon. Henceforth, absence of DJ-1 protein may trigger mechanisms which are not dependent upon α-Synucleinopathy. Interestingly, we found that ILCs are reduced in DJ-1^−/−^ mice. Thus, innate immunity is severely compromised such as high abundance of *Alistipes* and *Rikenella*. Previous studies suggested that *Alistipes sp* thrived in the absence of lipocalin 2 and IL-10^−/−^ mice and induced the intestinal inflammation and cancer progression in those animals (*70*). Moreover, NOD2 or RIP2 deficiency resulted in pro-inflammatory microenvironment that enhanced epithelial dysplasia following chemical injury and causes gut dysbiosis probably due to higher abundance of *Rikenella* bacterium in these mice (*75*). Interestingly, *Rikenella* was also shown to flourish in MyD88-deficient mice as these mice also lacks suitable innate immune system (*76*). Gender-specific differences found in the immune system and gut microbiome composition in males and females apparently fosters the expansion of *Alistipes, Rikenella* and *Porphyromonadaceae* in the absence of innate immune defence mechanism in male mice (*77*). These bacterial groups were linked with induction of weight loss, inflammation and DNA damage upon transfer of the male microbiota to germ-free female recipients (*77*). This study points out that it could be the case as DJ-1^−/−^ mice have lower numbers of ILCs and increased inflammation as well as higher abundance of *Alistipes* and *Rikenella*.

### Dysregulation of metabolites and neurodegeneration

The gut microbiome is a complex biological system and exhibits various tasks for the host including digestion, degradation of macromolecules, vitamin production and educating of the host innate and adaptive immune system (*78*). The gut microbiome composition affects the health of host *via* bacterial metabolites including SCFAs, TMAO and other metabolites, respectively (*66*). Bacterial metabolites appear to have diverse effects on metabolism and immune response and are considered as biomarkers for disease risk factor, whereas bacterial components cause an innate inflammatory response (*66*). Our metabolites data suggested that amino acids including valine, leucine, phenylalanine, alanine, tyrosine, and isoleucine were downregulated, whereas SCFAs including malonate, dimethyamine and Acetoin are upregulated in DJ-1^−/−^ mice. Defects in mitochondrial energy metabolism have been involved in the pathology of several neurodegenerative diseases (*79*). Metabolites generated during the bacterial metabolism and oxidation of the neurotransmitter dopamine (DA) are considered to damage the neurons of the basal ganglia (*79*). Infusion of metabolic inhibitor malonate into the striatum of mice or rats produced generation of DA nerve terminals and malonate induces a substantial increase in DA efflux in awake, behaving mice as quantified by in vivo microdialysis (*79*). Decreased SCFAs butyrate/propionate concentrations suggest loss of lactate utilizing bacterial strains (*80*). In PD patients fecal SCFAs were reduced compared with healthy controls (*31*). However, in DJ-1^−/−^ mice SCFAs were similar to those of WT mice. Furthermore, mitochondrial respiratory complex II (CII) is a protein complex located in the inner membrane of mitochondria and it forms part of the electron transport chain and is involved in succinate signalling and reactive oxygen species (ROS) (*81*). Malonate is a competitive inhibitor of the CII and reduces the cellular respiration, whereas succinate is rather drives the CII activity in macrophages (*82, 83*). Thus, metabolic stress induced by malonate, dimethyamine and acetoin could be involved in the neurodegenerative process in DJ-1^−/−^ mice potentially to be generated from bacterial metabolism or colon tissues.

The majority of the amino acids in the intestines derive from the metabolism of ingested dietary proteins, host tissue proteins or the conversion of other nitrogenous substances, whereas a small amount of amino acids is *de novo* synthesized by the gut bacteria (*84*). Amino acids, such as L-glutamine function as a double-edged sword for gut health as they can sponsor to the expression of pathogenic virulence genes as well as protect against disease (*84*). Most of the amino acids including valine, leucine, isoleucine and alanine were downregulated in the feces of DJ-1^−/−^ mice. Our results corroborate findings on PD patients samples derived from cerebrospinal fluid (CSF) for the amino acids including valine, leucine, isoleucine (*85*). PD patients might suffer from dysfunction of the transport of neutral and basic amino acids across the blood-brain barriers (*85*). Changes in amino acid metabolism in plasma/CSF of the patients, highlighting the role of altered amino acid metabolism and PD pathology (*86*). Decreased amino acid concentrations could be involved in loss of microbial proteases/peptide catabolism (*80*). Apparently, there is a switch from protein to fibre utilization under DJ-1 deficient conditions (*87*). PD patients who are not treated with levodopa or with dopamine agonists were reported to have higher CSF tyrosine and phenylalanine levels than those not treated with these drugs and also than controls (*85*). In this study, we found less tyrosine and phenylalanine in the feces of DJ-1^−/−^ mice compared with WT control group. Aromatic amino acids tyrosine and phenylalanine could be the most important amino acids as they are involved in L-DOPA synthesis by tyrosine hydroxylase (TH), which is further converted into dopamine, norepinephrine (noradrenaline), and epinephrine (adrenaline) (*88*). It is shown that 3-HPPA is formed through fermentation of tyrosine by *Clostridium* species (*61, 62*). Reduced levels of microbial tyrosine and phenylalanine point out that decreased production of dopamine precursors while enhanced levels of 3-HPPA which are able to cross into the brain might have a directive effect on dopamine synthesis e.g. as competitive inhibitor. However, further studies are warranted to understand the role of amino acid metabolism by gut bacteria in PD pathophysiology.

## Conclusion

Our study suggests that DJ-1^−/−^ mice present with gut dysbiosis, reduced innate lymphoid cells numbers or development, increased inflammation in the colon and feces, and dysregulated metabolites production already at an early disease stage (4 months) which could be toxic to colon tissues or neurons. Therefore, these disease markers could subsequently be explored for biomarker development in the PD patients.

## Material and Methods

### Animal breeding and ethical permission and materials collection

DJ-1^−/−^ mice were described earlier (*7*) and obtained from Prof. Tak W Mak, Canada. As described earlier, F1 progeny were backcrossed for seven generation to C57BL/6 mice and heterozygous animals (male and female) were used to set up the breeding to obtain homozygous for the targeted DJ-1 allele. Genotypes of mice was performed using PCR (WT *DJ-1* forward primer, TGC TGA AAC TCT GCC ATG TGA ACC; WT *DJ-1* reverse primer, CCT GCT TGC CGA ATA TCA T; and Neo, AGG TGA CAC TGC CAG TTG CTA GTC). PCR conditions were used as follow: 95°C for 30 sec, 64°C for 30 sec, and 72°C for 1 min (40 cycles). Age matched 3-4 months old DJ-1 deficient mice and control WT littermates from the same cohort were used for the feces collection for 16S rRNA microbiome analysis and stored at −80 °C until use. Some animals were sacrificed using CO_2_ methods and colons (3-4 cm long piece from at the junction of ceacum) and were collected in 4% PFA and snap frozen in liquid N_2_. All the animals were kept in the open cage in a standard environment mice facility. All the experiments were performed according to the EU Animals Scientific Procedures Act and the German law for the welfare of the animals. All the procedure and methods were approved by the local government authorities (Regierungspräsidium, Tübingen according to §4 animal welfare act on 20/07/2017) of the state of Baden-Württemberg, Germany.

### Bacterial DNA isolation from the feces

Frozen feces were weighted (50mg/sample) on dry ice and hammered to break into powder form and transferred into a 2.0 ml Eppendorf tube and kept on dry ice. Once all the samples were measured, all the weighted sample tubes were transferred to ice until the samples were thawed and added 1.0 ml of lysis buffer from QIAamp Fast DNA Stool Mini Kit (Cat no. #51604; Qiagen, Germany). All the procedures were followed as recommended by manufacturer’s guidelines for bacterial DNA isolation and DNA was dissolved in 100 μl instead of 200 μl DNA buffer.

### 16S rRNA sequencing

For 16S rRNA amplification, 12.5 ng of DNA was amplified using 0.2 μM of both forward primer (TCGTCGGCAGCGTCAGATGTGTATAAGAGACAGCCTACGGGNGGCWGCAG,
Metabion)
and reverse primer (GTCTCGTGGGCTCGGAGATGTGTATAAGAGACAGGACTACHVGGGTATCTAATC
C, Metabion).

KAPA HiFi HotStart Ready Mix (KK2601; KAPABiosystems) was used for the PCR amplification. PCR was performed using a first denaturation of 95°C for 3 minutes (min), followed by 25 cycles of amplification at 95°C for 30 s, 55°C for 30 s and 72°C for 30s, final elongation at 72°C for 5 min and the amplified DNA was stored at 4°C. DNA gel electrophoresis of all the samples was performed to verify the amplicon specificity.

Further, samples were then purified (Agencourt AMPure XP, Beckman Coulter) and PCR amplicons were indexed using Nextera XT index and KAPA HiFi HotStart ReadyMix. PCR was performed using a first denaturation of 95°C for 3 min, followed by 8 cycles of amplification at 95°C for 30 s, 55°C for 30 s and 72°C for 30, final elongation at 72°C for 5 min. Samples purified were then validated using BioAnalyzer (Bioanalyzer DNA 1000, Agilent) and 4 nM of each library pooled using unique indices before sequencing on a MiSeq (Illumina) and paired 300-bp reads.

### Sequence Analysis and Statistics

Available sequence data has been trimmed and filtered using SeqPurge (*89*). Trimming parameters demanded a minimum quality at 3’ end of q=35 (parameter qcut=35). Processed sequence data has been aligned using MALT (version 0.3.8; https://ab.inf.uni-tuebingen.de/software/malt) against the 16S database SILVA SSU Ref Nr 99 (https://www.arb-silva.de/documentation/release-128/) and classified using NCBI taxonomy. Alignment has been performed using semi-global alignment and a minimum sequence identity of 90% (parameter minPercentIdentity=90). Further analysis and visualization were performed using MEtaGenome Analyzer-Community Edition (MEGAN-CE) version 6.14.2, built 23 Jan 2019 (*42*) as describer earlier (*90*).

### Preparation of tissue for histological analysis

Colon samples prepared on ice were fixed for 24 h in 4% paraformaldehyde (PFA), stored at 4°C in 0.4% PFA for a maximum of 4 weeks prior embedding in paraffin. Fixed samples were then alcohol-dehydrated and embedded in paraffin. Samples were embedded in paraffin blocks using a tissue embedding station and stored at room temperature until use. Paraffin blocks containing colon tissues were cut into 7 μm thick sections using a microtome. Section were placed in 45°C water bath for flattening, collected on a glass slide, dried in an incubator at 50°C for 1-2 h and stored at room temperature.

### Immunohistochemistry

First slides were deparaffinize using autostained program 6 for a run for 51 minutes and slides were kept in TBS until antigen retrieval step. Antigen retrieval was done using sodium citrate buffer (1x) and citric acid (1x) method. Slides were boiled in the citric acid + sodium citrate buffer for 5 minutes for 3 times (3x) and slides were allowed to cool down for 15 minutes in TBS. After cooling slides were blocked of endogenous peroxidase and incubated for 20 minutes at room temperature (RT) and washed quickly 3x with TBS. Further slides were blocked for unspecific bindings using 5% normal goat serum in 0.3% Triton X-100 TBS and incubate for 1 hour at RT on slow rotation shaker. After incubation with unspecific binding, slides were washed for 3×3 minutes with TBS. Antibody staining was performed using α-Synuclein antibody (# 610786 BD Biosciences, Netherlands) for overnight (1:1000) dilution at 4 °C. Slides were washed next day with 0.025% Triton X-100 TBS for 3×5minutes. Secondary antibody was used (1:1000) dilution for 1 hour at RT. Further slides were stained with ABC complex-DAB for detection of primary antibody staining.

### Tissue lysate preparation for WB

Colon tissue were weighted frozen and lysed with 10 volumes of RIPA buffer (50 mM Tris, 150 mM NaCl, 1.0% NP-40, 0.5% sodium deoxycholate, 0.1% SDS, pH 8.0) supplemented with protease inhibitor (Complete; Roche Diagnostics). Colon tissues were homogenized for 1 minute using a disperser (T10 ULTRA-TURRAX; VWR) on ice. After the homogenization, samples were incubated for 30 min at 4°C and spun for 20 min at 12,000 g at 4 °C. Proteins lysate supernatants were supplemented with 10% glycerol before long storage at −80°C. Protein concentration was determined using BCA method (#23225; Thermofisher, Germany).

### Immunodetection

Samples were prepared by diluting protein lysates in PAGE buffer (0.2 M glycine, 25 mM Tris, 1% SDS), followed by a denaturation at 95°C for 10 min in loading buffer (80 mM Tris, 2% SDS, 5% 2-mercaptoethanol, 10% glycerol, 0.005% bromophenol blue, pH 6.8) and a short centrifugation 30 sec at 400 g. Proteins were separated by electrophoresis using 12% SDS-PAGE gel. Gels containing proteins were washed for 5 minutes in transfer buffer (0.2 M glycine, 25 mM Tris, 10–20% methanol) and transferred to membranes equilibrated in transfer buffer. Transfer was performed for 90 min at 80 V at 4°C on nitrocellulose membranes (88018, Life Technology). Immunoblot were washed 5 min in TBS buffer, fixed with 4% PFA for 1 hour (only for a-Syn detection) and blocked using 5% non-fat milk (Slim Fast) in TBS. Membranes were then washed twice 5 min in TBST and incubated with the primary antibody over night at 4°C (human and mouse a-syn: 610786 BD Biosciences; GAPDH: #5174, Cell Signaling and GFAP #MAB360, Merck Millipore, Germany). After incubation with the first antibody, membranes were washed four times (5 min each) with TBST. Membranes were then incubated for 75 min with the secondary antibody coupled to horseradish peroxidase (GE Healthcare). After four washing steps with TBST (5 min each), bands were visualized using the enhanced chemiluminescence method (ECL+; GE Healthcare). Light signal was detected using LI-COR Odyssey and were quantified using Odyssey software.

### Calprotectin ELISA and LEGENplex inflammatory cytokines

To measure the calprotectin/MRP 8/14 from the fecal samples, S100A8/S100A9 ELISA kit rat/mice (#K6936, Immundiagnostik AG, Germany) as well as LEGENplex™ (#740150 or #740446, Biolegend, Germany) mouse inflammation panel (13-plex) were used according to the manufacture’s guidelines. Fecal samples were measured (weight between 50 ± 5.0 mg) and dissolved in 500 μl of extraction buffer supplied by the kit, mixed by vortexing and then centrifuged for 10 minutes at 3000 g. Supernatant was taken and transferred to a new microcentrifuge tube and 100 μl of sample was used for measuring the protein for Calprotectin, however, for LEGENDplex inflammatory panel 25 μl of sample was used respectively. The data was analyzed using 4 parameters algorithm and the concentration of calprotectin/inflammatory cytokines was normalized with feces weight and data are presented in ng/g or pg/g.

### Flow cytometry

Splenocytes from WT and DJ-1-deficient mice were characterised by using surface and intracellular staining with relevant antibodies. In brief, splenocytes were collected and used for surface staining for dump-FITC lineage markers - CD3, CD5, CD3, CD11b, CD11c, F4/80, Gr-1, B220 or CD19, and Ter119 and CD45-PE (eBioscience, Germany) washed with PBS. Cells were fixed with Foxp3 fixation/permeabilization buffer (eBioscience, Germany) for intracellular staining and incubated for 30 minutes. After incubation, cells were washed with 1× permeabilization buffer, exposed to added intracellular monoclonal antibodies for RoRgt-PerCP-Cy5.5, Eomes-APC, Tbet-PerCP-Cy5.5 and GATA3-APC and incubated for an additional 45 minutes. Cells were washed again with permeabilization buffer and PBS was added to acquire the cells on a flow cytometer (FACS-calibur™ from Becton Dickinson; Heidelberg, Germany).

### Feces sample prep for the metabolite detection using ^1^H-NMR

For metabolite extraction, 50 mg of deep-frozen feces sample were transferred into 2 mL AFA glass tubes (Covaris Inc, Woburn, USA) and mixed with 400 μL of ultrapure methanol and 800 μL of MTBE (solvent grade). The mixture was manually dispersed with a disposable plastic spatula, then vortexed and transferred to a focused ultrasonicator (Covaris E220evolution, Woburn, USA). Feces metabolites were extracted with a 5 min lasting ultrasonication program in a degassed water bath at 7° C. After extraction, the metabolite suspension was separated into a polar and lipid phase by adding 400 μL of ultrapure water. In order to remove and remaining solids from the samples, the glass tubes were centrifuged for 5 min at 4000 rpm. 700 μL of each phase were then transferred to a fresh 1.5 mL Eppendorf cup. The polar phase was subject to a 2nd centrifugation step for 5 min at 12,000 rpm and 600 μL of the supernatant were transferred to a new 1.5 mL Eppendorf cup and evaporated to dryness over night with a vacuum concentrator (Eppendorf Speedvac).

### ^1^H-NMR for metabolites and data analysis

For NMR analysis, the dried pellets were resuspended with 60 μL of deuterated phosphate buffer (200 mM K2HPO4, 200 μM NaN3, pH 7.4) containing 1 mM of the internal standard TSP (trimethylsilylpropanoic acid). In order for a maximum dissolution, the plastic tubes were quickly sonicated and then centrifuged 5 min at 14,000 rpm. 50 μL of the supernatant were transferred with gel loading pipette tips into 1.7 mm NMR tubes (Bruker BioSpin, Karlsruhe, Germany) and a 96 well rack placed into the cooled (4° C) NMR autosampler.

Spectra were recorded on a 600 MHz ultra-shielded NMR spectrometer (Avance III, Bruker BioSpin GmbH) equipped with a triple resonance (1H, 13C, 31P) 1.7 mm room temperature probe at 298 K. For optimum water suppression and shim adjustment a quick simple ZG experiment was performed followed by a 1h lasting CPMG (Carr-Purcell-Meiboom-Gill) experiment in order to suppress residual background signals from macromolecules such as bilirubin (time domain = 64k points, sweep width = 20 ppm, 512 scans). The recorded free induction decays (FIDs) were fourier-transformed and spectra properly phase- and baseline corrected. Metabolite annotation and quantification was performed with ChenomX NMR Suite 8.3 and statistical analysis with MetaboAnalyst 4.0.

The phenol derivative 3-(3-Hydroxyphenyl) propanoic acid 3-HPPA was identified by selective TOCSY experiments followed by purchasing possibly fitting reference standards of different phenols with substituted groups in the meta position. Reference spectra of those were recorded in the used feces extract phosphate buffer and by applying spiking experiments.

### Statistical analysis

MEGAN-CE (version 6.14.2, built 23 Jan 2019) and MicrobiomeAnalyst were used for data acquisition and analysis. GraphPad and Inkscape were used for the final figure preparation. One-way ANOVA or Student’s t-test was used for statistical analysis using GraphPad wherever it was appropriate and described in the figure legend. Data shown in either Means±SD or SEM. The p value (≤0.05) considered significant. Metabolite concentrations from ^1^H-NMR analysis were exported as comma separated value spreadsheet file to MetabAnalyst, normalized with PQN (probabilistic quantile normalization) and range scaled and then analyzed with student’s t-test, oPLSDA and VIP analysis.

## Supporting information

Suppl. File Singh Y et al DJ1

## Fig. legends

Suppl. Fig. 1: KEGG functional gene pathway analysis from 16S rRNA.

Suppl. Fig. 2: Inflammatory cytokine production in the feces of DJ-1^−/−^ and WT mice.

Suppl. Fig. 2: NMR Metabolites Heatmap and metabolites clustering in WT and DJ-1^−/−^ feces.

Suppl. Fig. 4: Example of ^1^H-NMR spectra of DJ-5 and recorded reference standard spectrum for 3-HPPA

Suppl. Fig. 5: α-Syn expression by IHC and Immunoblotting. (a) A representative IHCs colon pictures from WT (n=3) and DJ-1^−/−^ (n=3) mice. Each mouse is shown and marked with #1, #2 and #3 and staining of α-Syn is shown with arrow heads in the pictures. (b) Bar diagram (mean±SEM) data shows similar expression in WT (n=6) and DJ-1^−/−^ (n=6).

## Acknowledgements

This is an EU Joint Programme - Neurodegenerative Disease Research (JPND) project. The project JPCOFUND_FP-829-047 aSynProtec is partly supported through the funding organization Deutschland, Bundesministerium für Bildung und Forschung (BMBF, FKZ). This research was supported by the Deutsche Forschungsgemeinschaft, DFG through the funding of the NGS Competent Centre Tübingen (NCCT-DGF). We thank Prof Bernd Pichler for allowing to have access to the ^1^H-NMR for metabolic studies. Funders have no role in the study design and data analysis. We acknowledge support by Deutsche Forschungsgemeinschaft and Open Access Publishing Fund of University of Tübingen.

## Ethics statement

All animal experimental protocols used in this study is strictly adhered to the international standards for the care and use of laboratory animals and were approved by the local Animal Welfare and Ethics committee of the Country Commission Tübingen, state of Baden-Württemberg, Germany (§4 animal welfare act on 20/07/2017).

## Author’s role

YS: Study design, performed the research and managed the overall project, involved in entire study, analysed the data, made the figures and wrote the manuscript

CT: Study design, performed the metabolites study, data analysis, made the figures, wrote the manuscript

AD, JM: Helped with 16S rRNA data analysis bioinformatics meta data analysis and made the figures

MA, MSS: study design, performed the research, helped with animal experiments

LZ: Helped with the identification of 3-HPPA, did reference standards and spiking experiments

LP: performed the animal experiments

MF, MF: Provided/tools provided for the study, performed and help with the animal experiments

VP: Helped with study design and data analysis

DSP, TWM: Materials/tools provided for the study

JSF: Helped with study design and discussion of the 16S rRNA data

NC: Study design, substantial discussion for 16S rRNA data

MSS, DW, SYB, FL, OR: Study design, funding generation, wrote the manuscript

All authors read the manuscript and approved to be co-authors on the manuscript and have substantial contribution in the manuscript.

