## Supplementary figures and images for "DJ-1 (Park7) affects the gut microbiome, metabolites and development of Innate Lymphoid cells (ILCs)"

### Suppl. File Singh Y et al DJ1

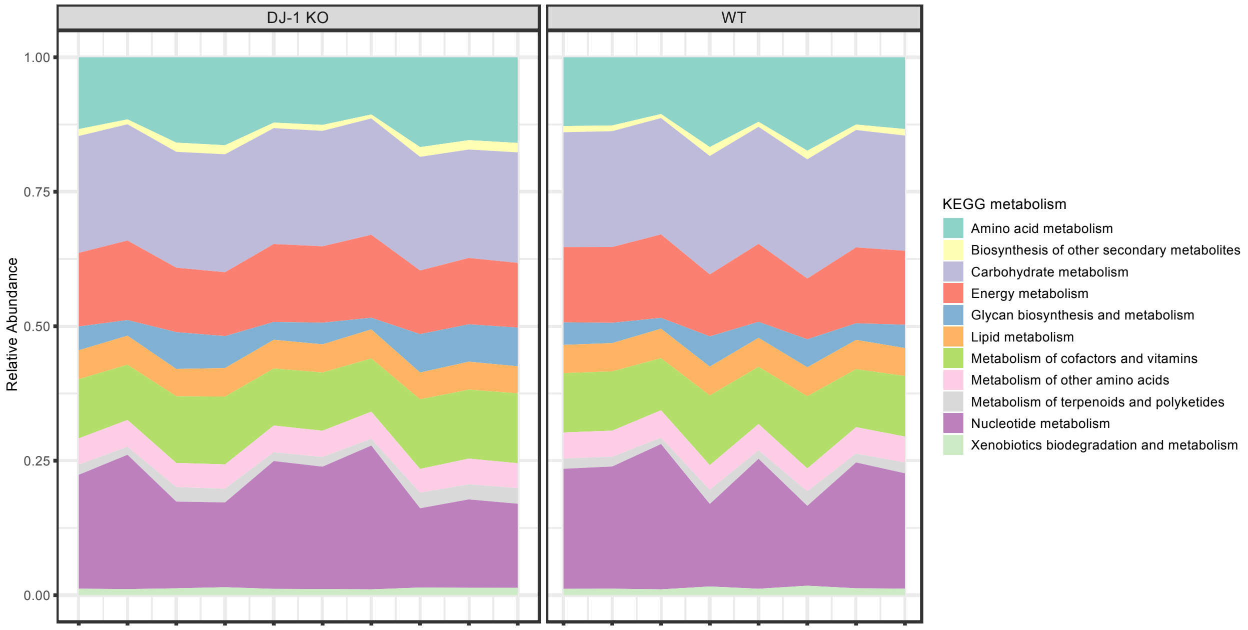

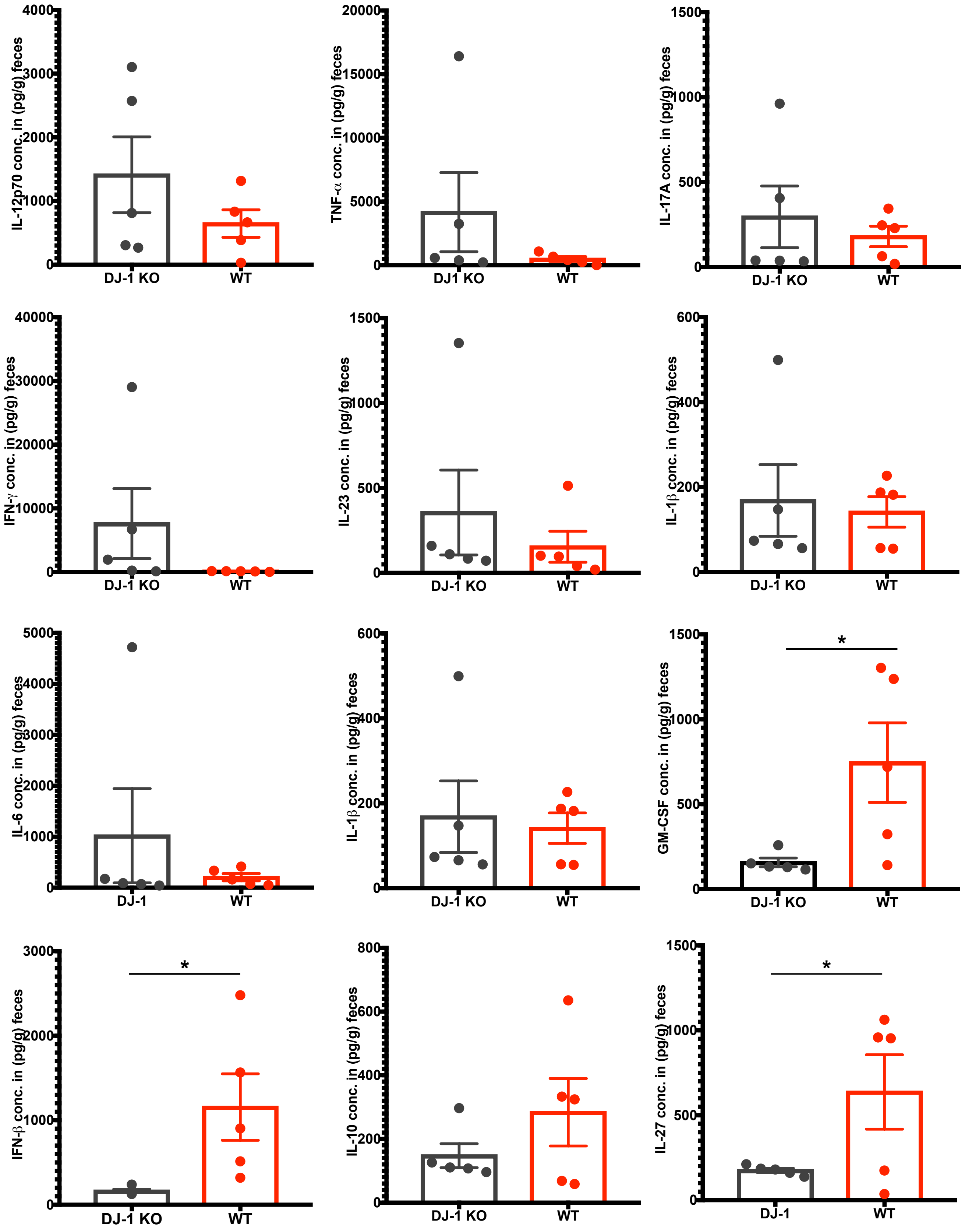

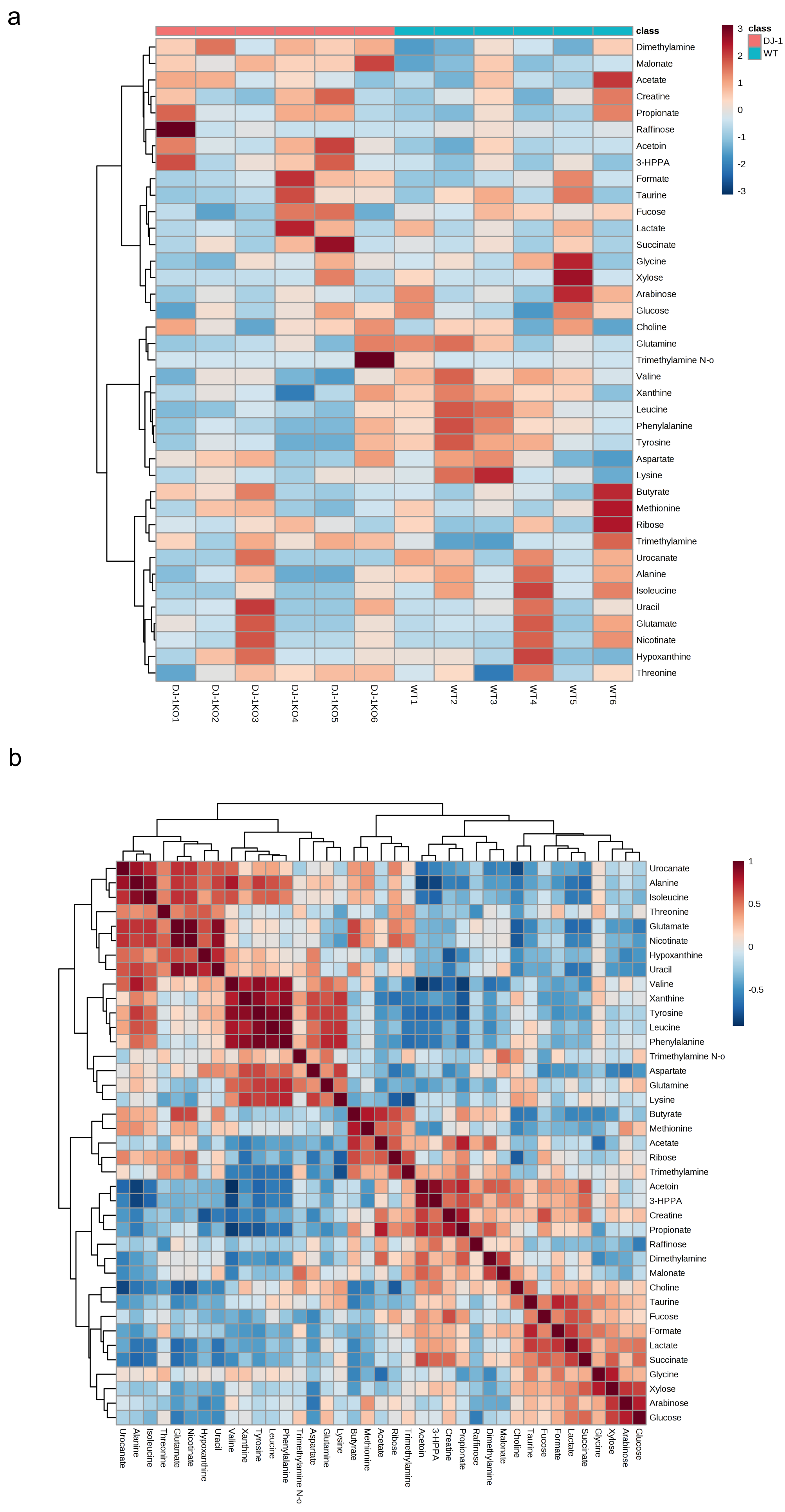

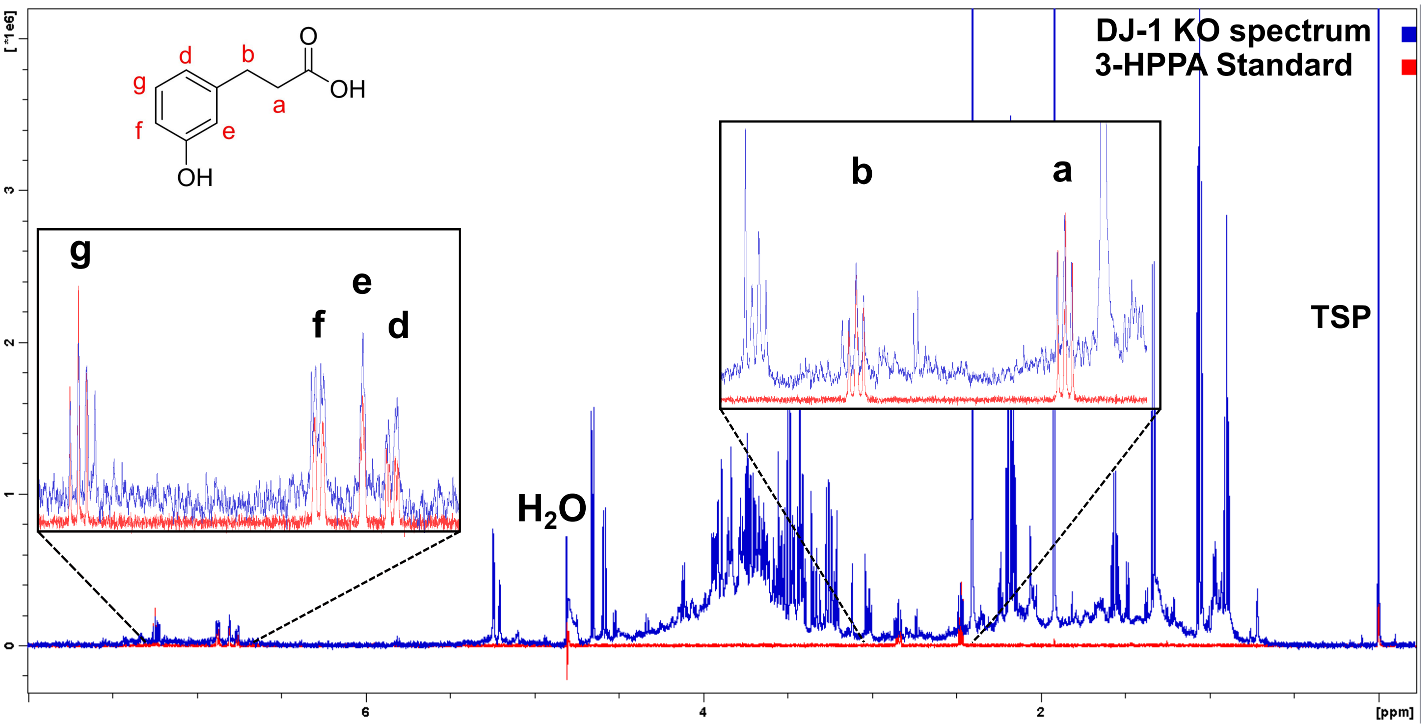


Suppl. Fig. 4

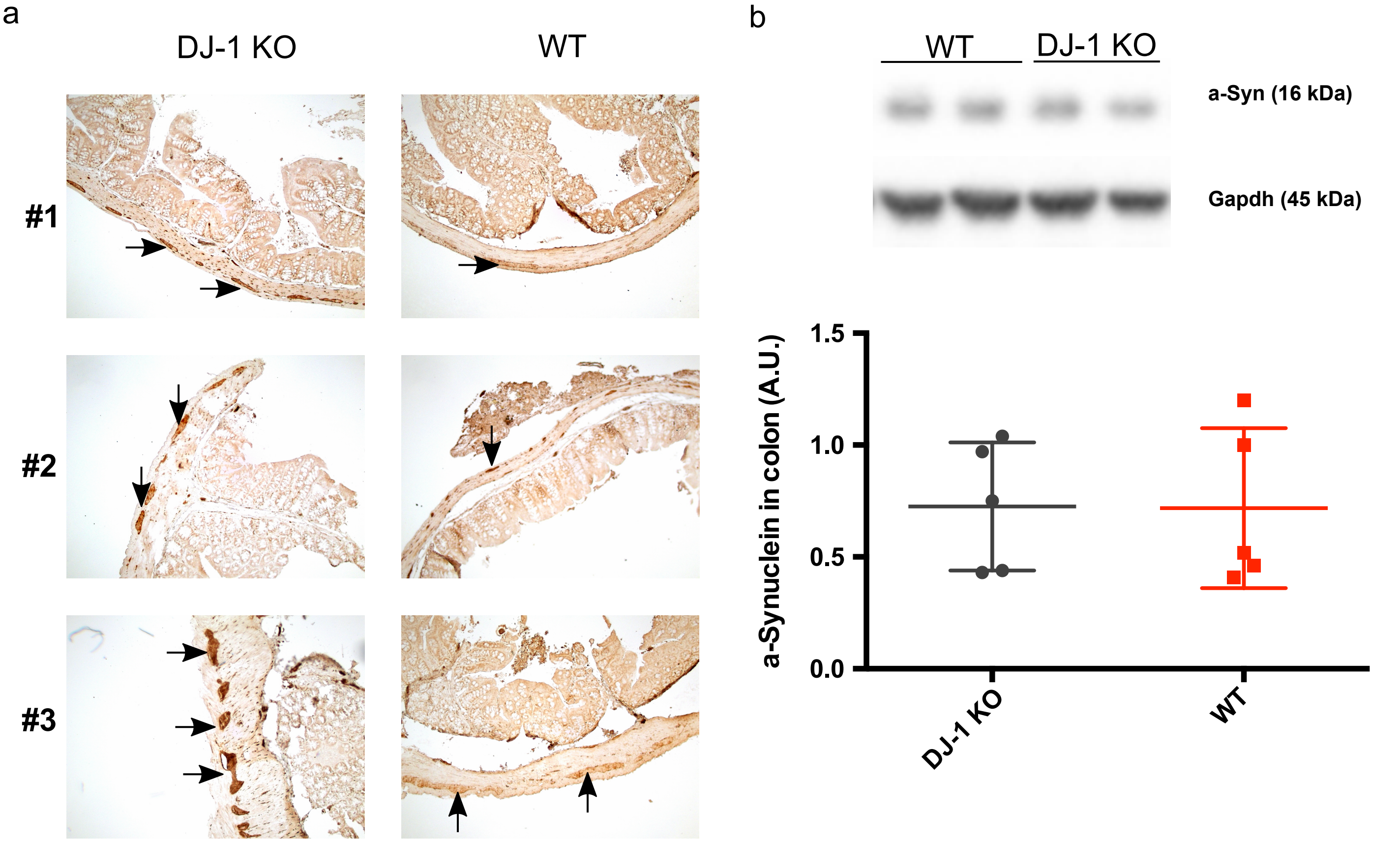
